# Differential contribution of P73^+^ Cajal-Retzius cells and Reelin to cortical morphogenesis

**DOI:** 10.1101/2024.10.15.618167

**Authors:** Vicente Elorriaga, Benoît Bouloudi, Yoann Saillour, Juliette S Morel, Elodie Delberghe, Patrick Azzam, Matthieu X Moreau, Rolf Stottmann, Nadia Bahi-Buisson, Alessandra Pierani, Nathalie Spassky, Frédéric Causeret

## Abstract

Cajal-Retzius cells (CRs) are a peculiar neuronal type within the developing mammalian cerebral cortex. One of their best documented feature is the robust secretion of Reln, a glycoprotein essential for the establishment of cortical layers through the control of radial migration of glutamatergic neurons. We previously identified *Gmnc* as a critical fate determinant for P73^+^ CRs subtypes from the hem, septum and thalamic eminence. In *Gmnc^-/-^* mutants, P73^+^ CRs are initially produced, cover the telencephalic vesicle but undergo massive apoptosis resulting in their complete depletion at mid-corticogenesis. Here we investigated the consequence of such a CRs depletion on dorsal cortex lamination and hippocampal morphogenesis. We found preplate splitting occurs normally in *Gmnc^-/-^* mutants but is followed by defective radial migration arrest in the dorsal cortex, altered cellular organization in the lateral cortex, aberrant hippocampal progenitor proliferation resulting in abnormal CA1 folding and lack of vasculature development in the hippocampal fissure. We then performed conditional *Reln* deletion in P73^+^ CRs to evaluate its relative contribution and found that only radial migration defects were recapitulated. We concluded that at mid-corticogenesis, CRs-derived Reln is required for radial migration arrest and additionally identified Reln-independent functions for CRs in the control of hippocampal progenitor proliferation and vessel remodelling.

## Introduction

Development of the mammalian cerebral cortex relies on the precise coordination of progenitor patterning and proliferation, specification of cellular identities and neuronal migration, that ultimately enable the establishment of mature functional circuits. Cajal-Retzius cells (CRs) are key players in these processes (Causeret et al., 2021; Elorriaga et al., 2023). CRs are among the earliest-born neurons of the cerebral cortex, mostly between E10.5 and E11.5 in mice (Hevner et al., 2003; Takiguchi-Hayashi et al., 2004), and originate from focal sources at the border of the developing pallium: the cortical hem dorso-medially (Meyer et al., 2002; Takiguchi-Hayashi et al., 2004), the pallial septum rostro-medially (Bielle et al., 2005), the thalamic eminence (ThE) caudo-medially (Ruiz-Reig et al., 2017; Tissir et al., 2009) and the ventral pallium (VP, also referred to as pallial-subpallial boundary, PSB) laterally (Bielle et al., 2005). Upon production, CRs migrate tangentially in the marginal zone and distribute over the entire telencephalic vesicle. At postnatal stages, CRs undergo apoptosis and almost completely disappear from the dorsal cortex whereas a fraction estimated to 15% survives until adulthood in the hippocampus (Anstötz et al., 2016; Chowdhury et al., 2010; Ledonne et al., 2016). A precise control of CRs presence and demise was shown to be essential for the establishment of cortical and hippocampal circuits (de Frutos et al., 2016; Genescu et al., 2022; Glærum et al., 2024; Riva et al., 2019; Riva et al., 2023).

CRs diversity characterized by single-cell transcriptomics primarily distinguishes medial populations (hem-, septum-and ThE-derived) that share the expression of a particular gene module exemplified by *Trp73* (encoding the transcription factor p73) (Moreau et al., 2021). By contrast, the lack of specific marker to distinguish VP-derived CRs from their p73^+^ counterparts so far hindered their direct visualization. Despite these differences, all CRs share a set of features, including expression of the transcription factors Tbr1 (reflecting their glutamatergic identity) and Nhlh2, as well as the secreted protein Reelin (Reln) (Hevner et al., 2003; Moreau et al., 2023; Ogawa et al., 1995).

Reln is an extracellular matrix glycoprotein strictly required to establish the inside-out layered pattern of the neocortex and the radial organization of the hippocampus (Boyle et al., 2011; Caviness and Sidman, 1973; D’Arcangelo et al., 1995). However, the precise function of Reln remains elusive, as it was both proposed to promote radial migration and act as a stop signal, pointing to the idea that Reln may exert distinct functions at different phases of the radial migration process (Dulabon et al., 2000; Jossin et al., 2004; Kubo et al., 2010; Sekine et al., 2014; Zhao and Frotscher, 2010). In addition, ectopic Reln expression in the ventricular zone of *Reeler* mutants was shown to rescue preplate splitting but not lamination (Magdaleno et al., 2002), indicating these two processes differentially rely on the amounts and localization of Reln.

The specific contribution of CRs to corticogenesis was previously investigated using genetic ablation paradigms. Ablation of septum-derived CRs (*Emx1^Cre^;Dbx1^DTA^*) was shown to result in apparently preserved lamination in the depleted region, correlating with only a temporary decrease in Reln before other CRs populations repopulate the depleted areas (Griveau et al., 2010). Hem-ablated mutants (*Emx1^Cre^;Wnt3a^DTA^*), that lack most neocortical CRs, were surprisingly reported to display only a moderate disruption of lamination, leading to the proposal that CRs-derived Reln is in part dispensable for layer formation (Yoshida et al., 2006). Of note, the absence of hippocampus in hem-ablated mutants precluded from analysing the impact of CRs loss on hippocampal morphogenesis. Additional models of CRs depletion are provided by constitutive or isoform-specific p73 mutants (Amelio et al., 2020; Medina-Bolívar et al., 2014; Meyer et al., 2004; Meyer et al., 2019; Tissir et al., 2009). Although a range of hippocampal and cortical morphogenesis defects were identified in these animals, only a few cell-type specific molecular markers were assessed, resulting in an incomplete understanding of the consequences of CRs loss at embryonic stages and pointing to the need for additional molecular characterization.

We recently pinpointed the molecular mechanisms involved in the fate specification of P73^+^ CRs from the hem, septum and ThE, through the repurposing of a gene regulatory network previously known to control the cellular process of multiciliogenesis (Moreau et al., 2023). Genetic disruption of this regulatory network by targeting *Gmnc*, the most upstream gene in the cascade, results in the induction of apoptotic signalling by P73^+^ CRs from E12, their massive death at E13 and near complete elimination at E14. To date, the impact of such a massive CRs loss after a short period of normal presence on cortical and hippocampal development remains unknown. Here, we first characterized the extent and dynamics of CRs and Reln loss in *Gmnc^-/-^*embryos. We then compared the phenotype of *Gmnc* mutants as a model of severe CRs depletion without hem alteration, with conditional *Reln* deletion in P73^+^ CRs subtypes to better characterize the relative contribution of CRs and Reln to cortical morphogenesis. We found that at mid-corticogenesis, CRs-derived Reln contribute to radial migration arrest in the dorsal cortex and additionally demonstrate that hippocampal CRs impact on progenitor proliferation and vasculature development most likely in a Reln-independent manner.

## Material and Methods

### Animals

The following mouse lines were used and maintained on a C57BL/6J background: *Gmnc^-/-^* (*Gmnc^tm1.1Strc^*) (Terré et al., 2016) *PGK^Cre^* (*Tg(Pgk1-cre)1Lni*) (Lallemand et al., 1998), *ΔNp73^Cre^* (*Trp73^tm1(cre)Agof^)* (Tissir et al., 2009), *Wnt3a^Cre^* (*Wnt3a^tm1(cre)Eag^*) (Yoshida et al., 2006), *Rosa26^tdTomato^* (*Gt(ROSA)26Sor^tm9(CAG-tdTomato)Hze^*) (Madisen et al., 2010) and *Reln^flox^* (*Reln^tm1c(KOMP)Mbp^*) (Cionni et al., 2016). All animals were handled in strict accordance with good animal practice, as defined by the national animal welfare bodies, and all mouse work was approved by either the French Ministry of Higher Education, Research and Innovation as well as the Animal Experimentation Ethical Committee of Université Paris Cité (CEEA-34, licence numbers: 2020012318201928 and 2018020717269338).

### scRNAseq

The forebrain of four *Gmnc^-/-^* P0 animals originating from two distinct litters and four wild-type littermates were collected and maintained in ice-cold Hank’s balanced salt solution (HBSS). The region encompassing the lateral and dorsal cortex as well as the hippocampus (Fig. 5A) was dissected on both hemispheres and samples were pooled according to their genotype. Cell dissociation was achieved using the Neural Tissue Dissociation Kit (P) (Miltenyi Biotec) and a gentleMACS Octo Dissociator following the manufacturer’s instructions. Cells were loaded on a 10X Genomics Chromium Controller and two Next GEM Single Cell 3’ v3.1 libraries were produced. Sequencing was performed on a NovaSeq 6000 sequencer (Illumina) and a SP flow cell for a total depth of 800 million reads. Reads were aligned to the mm10 reference genome using Cell Ranger v7.1.0.

Subsequent analyses were performed using the Seurat v5.0.1 (Hao et al., 2024) package under R v4.1.1. We filtered-out cells with less than 800 genes, less than 3000 transcripts or more than 10% mitochondrial reads. Genes expressed in less than 3 cells were also excluded from the count matrix. Predicted doublets were identified using Scrublet (Wolock et al., 2019) and removed. The wild-type and mutant datasets were merged without integration and dimensionality reduction was achieved using the SPRING tool (Weinreb et al., 2018).

### Data availability

Raw and processed data, including a Seurat object corresponding to Fig 6B, can be retrieved from the GEO database (accession number GSE276037). Comprehensive and annotated R codes used for quality control, analysis and figure layout can be found at https://fcauseret.github.io/P0_GmncKO/.

### FlashTag and EdU injections

EdU was delivered by intraperitoneal injection (100µL at 1mg/mL) in E12 pregnant females. At E14, the same animals were injected with 100µg/kg buprenorphine 30 min prior to anaesthesia with Isoflurane (4% induction, 2% during the surgery) and subjected to abdominal incision to expose the uterine horns. CellTrace™ CFSE (or FlashTag, Thermo Fisher Scientific) was reconstituted at 5mM in DMSO and mixed with 10% Fast Green. Approximatively 1µL was injected into the lateral ventricles of embryos with a glass capillary. Uterine horns were repositioned into the abdominal cavity, and the abdominal wall and skin were sutured. Post-surgery analgesia was ensured using 5mg/kg Ketoprophen twice a day for 72h. Embryos were harvested at E18.

### Tissue processing

Embryonic and postnatal tissue were collected in ice-cold HBSS, immediately fixed by immersion in 4% paraformaldehyde, 0.12 M phosphate buffer pH 7.4 (PB) for 4h at 4°C, cryoprotected by overnight incubation in 10% sucrose in PB at 4°C, embedded in 7.5% gelatin, 10% sucrose in PB, and frozen by immersion in isopentane cooled at −55°C with dry ice. 20 µm coronal sections were obtained with a Leica CM3050 cryostat and collected on Superfrost Plus slides (Menzell-Glasser).

### Immunostaining

The following primary antibodies were used: goat anti-Brn2 (POU3F2, Abcam ab101726 1:1000), rat anti-CTIP2 (BCL11B, Abcam ab18465 1:600), rabbit anti-FOXG1 (Abcam ab18259 1:2000), rabbit anti-Laminin (Sigma-Aldrich L9393 1:600), goat anti-Neuropilin-1 (R&D Systems AF566 1:800), rabbit anti-p73 (Cell signaling 14620 1:250), goat anti-Nurr1 (NR4A2, R&D Systems AF2156 1:200), goat anti-Prox1 (R&D Systems AF2727 1:1000), goat anti-Reelin (R&D Systems AF3820 1:2000), rabbit anti-TBR1 (Abcam ab31940 1:1000).

Secondary antibodies directed against goat, rabbit, rat or mouse IgG and coupled to Alexa-488, Cy3 or Cy5 were obtained from Jackson ImmunoResearch. DAPI (1 µg/ml) was used for nuclear staining. Slides were mounted in Fluoromount-G (Thermo Fisher Scientific).

EdU detection was performed using the Click-iT™ EdU Alexa Fluor™ 647 Imaging Kit (Thermo Fisher Scientific) according to the manufacturer’s instructions.

### In situ hybridization

For each gene of interest, a 500-800 bp DNA fragment was amplified by PCR from a mouse embryonic brain cDNA library using Phusion polymerase (Thermo Fisher Scientific). Primers for *Nr4a2*, *Tbr1* and *Zbtb20* are described in (Moreau et al., 2021), and those for *Nhlh2* in (Moreau et al., 2023). *Gad2* was amplified using the following forward and reverse primers: TCTTTTCTCCTGGTGGCG and TTGAGAGGCGGCTCATTC. In all cases, the promoter sequence of the T7 RNA polymerase (GGTAATACGACTCACTATAGGG) was added in 5′ of the reverse primer. Alternatively, for *Reln*, and *Rorb*, a plasmid containing part of the cDNA was linearized by enzymatic restriction. Antisense digoxigenin-labelled RNA probes were then synthetized by *in vitro* transcription using T7 RNA polymerase (New England Biolabs) and digRNA labelling mix (Roche). *In situ* hybridization was carried out as previously described (Schaeren-Wiemers and Gerfin-Moser, 1993) using a buffer composed of 50% formamide, 5X SSC, 1X Denhardt’s, 10% dextran sulfate, 0.5 mg/mL yeast RNA, 0.25 mg/mL herring sperm DNA. Probes were detected using an anti-digoxigenin antibody coupled to alkaline phosphatase (Roche) and NBT/BCIP (Roche) as substrates. Slides were mounted in Mowiol.

### Image acquisition

Images were acquired using a Hamamatsu Nanozoomer 2.0 slide scanner with a 20× objective and a Leica SP8 confocal microscope with a 40× objective. Images were analysed using ImageJ software and figures panels assembled with Adobe Photoshop.

### Quantifications

Images were analysed using Fiji (https://fiji.sc/). Tbr2^+^ cell numbers (Fig 1F) were obtained by manually counting at least 3 sections from n= 2 animals per stage and per genotype. Cell numbers positive for p73 or Tomato (Fig 2E, 7D, 7E) were automatically counted on confocal z-stacks using the Cellpose wrapper (*Cellpose Advanced*, version 2.0 (Stringer et al., 2021) of the PTBIOP plugin (https://biop.epfl.ch/Fiji-Update/). n= 3 animals per genotype were considered. FlashTag and EdU-positive cells (Fig 3F) were also detected automatically in representative columns of 400 µm width and their radial position was normalized according to the thickness of the cortical plate (from subplate to MZ). 6 sections from 3 animals were considered for each genotype, over 3200 cells were counted with no less than 680 cells per condition. Plots were obtained using the geom_density() function from the R package ggplot2 with default parameters. Similarly, the density of Tomato^+^ cells along the medio-lateral axis of the cortical MZ (Fig 7E, 7F) was computed by measuring the relative coordinates of over 2200 cells collected from 3 to 5 sections from n=3 controls and 3 mutants, and no less than 70 cells per section. Plots were obtained using the geom_density() function from the R package ggplot2 with default parameters. Reln immunostaining intensity measurement (Fig 2C and 7G) were performed using the *Plot profile* function of Fiji on a 55µm-thick segmented line starting from the distal tip of the hippocampal fissure (HF) and following the MZ up to the piriform cortex in 9 sections from 3 animals for each genotype. Data were normalized by the total length of the segmented line. Plots were obtained using the geom_smooth() function from the R package ggplot2 with default parameters. MZ and cortical plate thickness (Fig 3B) were measured manually using DAPI staining. We considered the somatosensory cortex of n = 3 animals for each genotype. Per animal, 2 images were taken in each brain hemispheres of 2 anatomically equivalent sections. EdU^+^ cells in the hippocampus (Fig 4D) were manually counted in a 200µm wide radial column, separating the *stratum oriens* and *stratum pyramidale* layers using DAPI staining. n = 3 animals per genotype were considered. In mutants, we counted two adjacent columns corresponding to a folded and a normal region. In controls, two adjacent normal columns were counted per animal.

**Figure 1.**
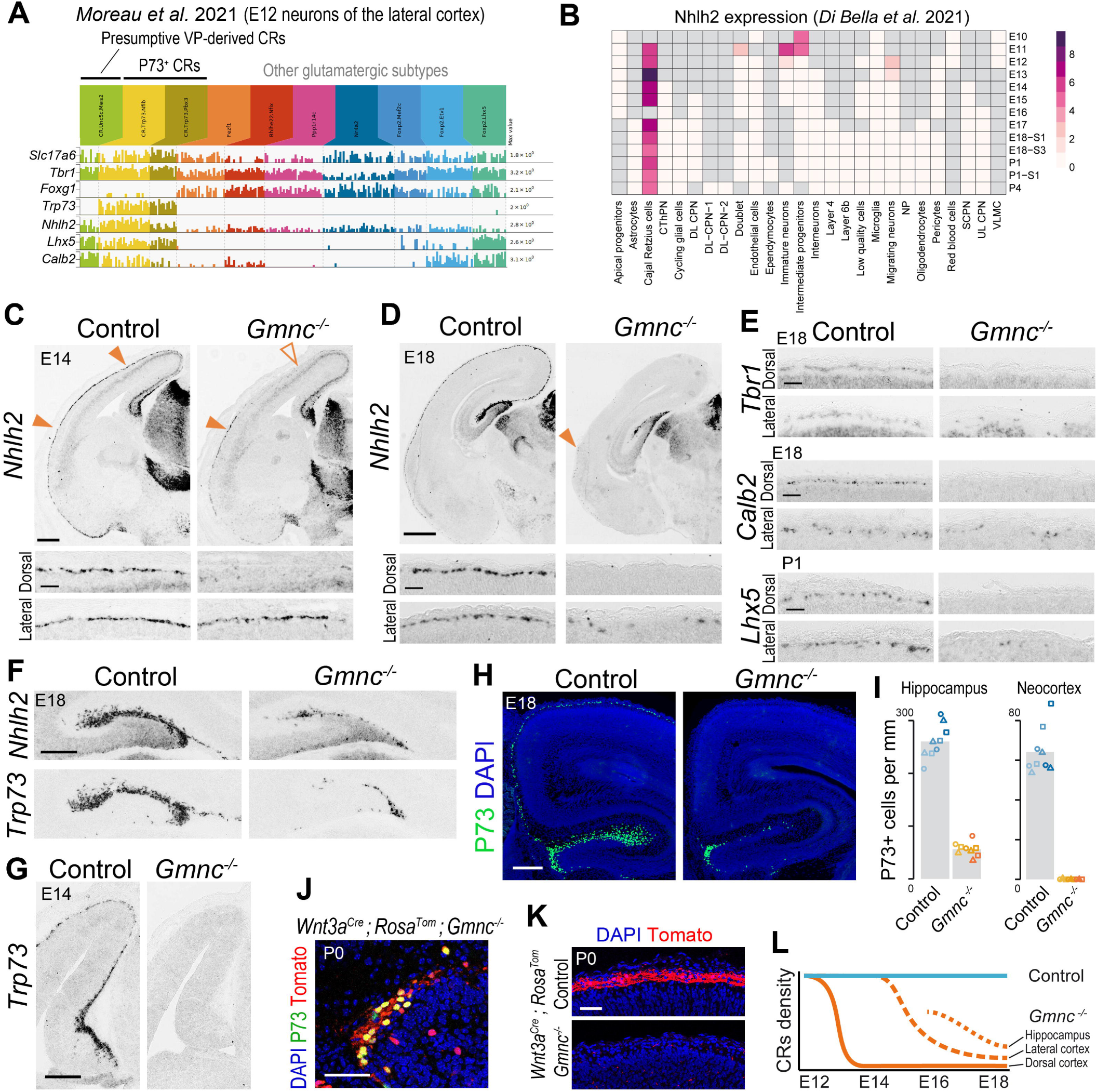
CRs depletion in *Gmnc* mutants. (A) Gene expression in E12 CRs subtypes and other glutamatergic neurons from the dorsal and lateral cortex (extracted from https://apps.institutimagine.org/mouse_pallium/). (B) Expression of *Nhlh2* per cell type and stage in scRNAseq data from the somatosensory cortex. Grey squares indicate no cells were sampled. Note that *Nhlh2* expression is restricted to CRs except at early stages where it is also detected in intermediate progenitors and immature neurons. (C, D) *In situ* hybridization for *Nhlh2* on coronal sections of the cerebral cortex from E14 (C) and E18 (D) control and *Gmnc^-/-^*embryos. The presence/absence of *Nhlh2*^+^ cells in the MZ is indicated by filled/empty arrowheads, respectively. (E) High magnification of the dorsal or lateral cortex MZ after *in situ* hybridization for *Tbr1* and *Calb2* at E18, and *Lhx5* at P1. (F) *In situ* hybridization for *Nhlh2* and *Trp73* on coronal sections of the E18 hippocampus. (G) *In situ* hybridization for *Trp73* on coronal sections of the E14 dorsomedial cortex. (H) Immunostaining for P73 on coronal sections of the E18 hippocampus and dorsomedial cortex. (I) Quantification of the density of P73^+^ cells in the hippocampal and neocortical MZ of E18 control and *Gmnc^-/-^*embryos. Each dot corresponds to one measurement, 3 animals (color-coded) and 3 rostro-caudal levels (shape) were considered. (J) Immunostaining for P73, and Tomato in the hippocampus at P0 following genetic tracing of hem derivatives in a *Gmnc^-/-^* background, showing residual hem-derived CRs in mutants. (K) Immunostaining for Tomato and DAPI in the dorsal cortex at P0 following genetic tracing of hem derivatives in either control or *Gmnc^-/-^* background, showing the complete absence of Tomato^+^ cells in the mutant neocortical MZ. (L) Schematic representation of the temporal dynamics of CRs depletion in *Gmnc^-/-^*mutants. Scale bars: 200µm in C, F, G, H, 500µm in D, 50µm in high magnification panels in C, D, E and in J, K.

**Figure 2.**
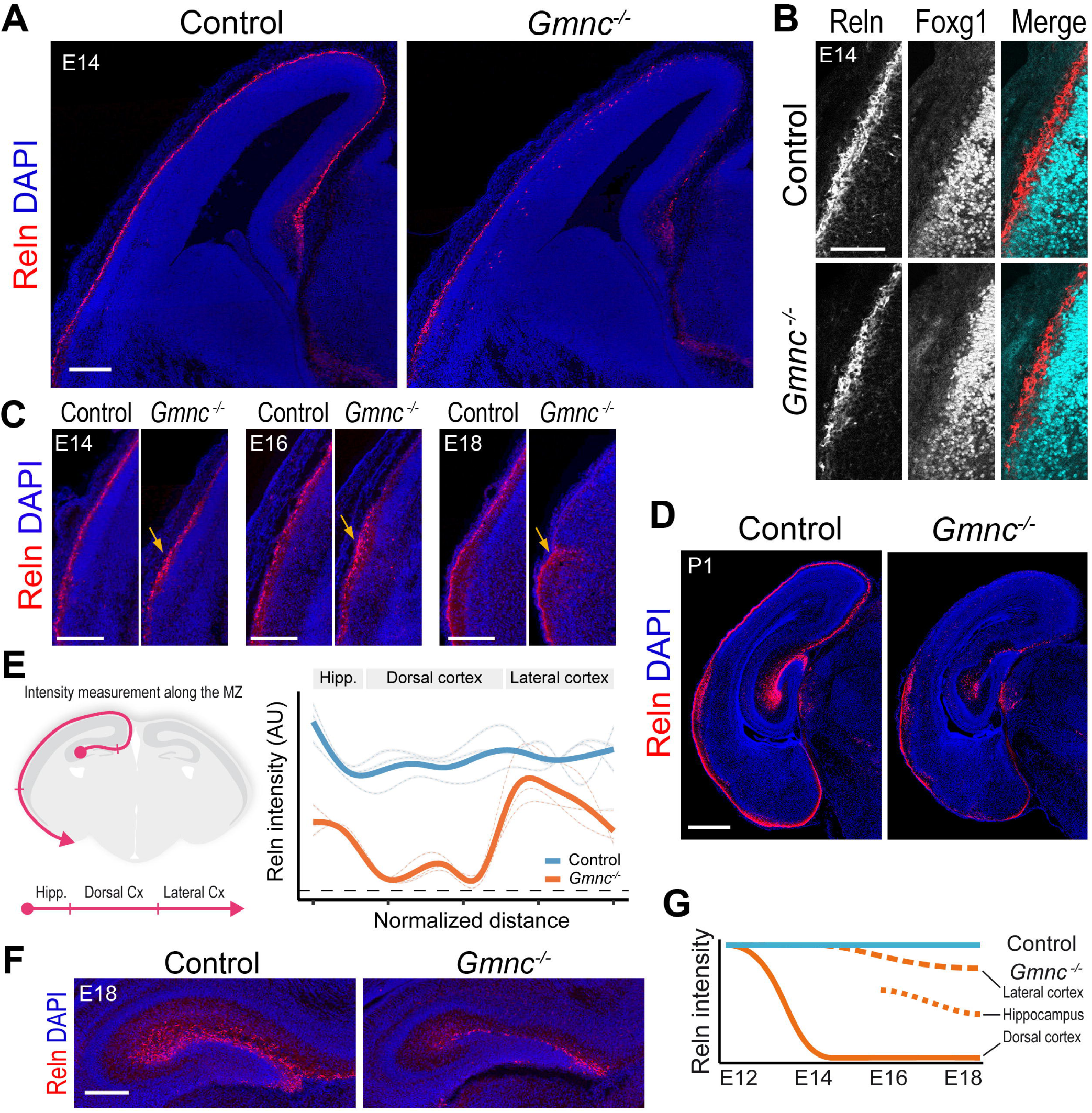
Reln depletion in *Gmnc* mutants. (A) Immunostaining for Reln on coronal sections of the cortex at E14. (B) Immunostaining for Reln and Foxg1 on the lateral cortex at E14. (C) Immunostaining for Reln in the lateral cortex at E14, E16 and E18. The arrow points to the progressive formation of a boundary between lateral and dorsal cortex in mutants. (D) Immunostaining for Reln on coronal sections of the cortex and hippocampus at P1 in control *Gmnc^-/-^* mutant. (E) Quantification of the Reln staining intensity in the MZ (arbitrary units) along the medio-lateral axis. The dashed lines represent the 3 animals for each genotype. (F) Immunostaining for Reln in the E18 hippocampus of control and *Gmnc^-/-^* mice. (G) Schematic representation of the temporal dynamics of Reln depletion in *Gmnc^-/-^* mutants. Scale bars: 200µm in A, B, C, F. 500µm in D.

**Figure 3.**
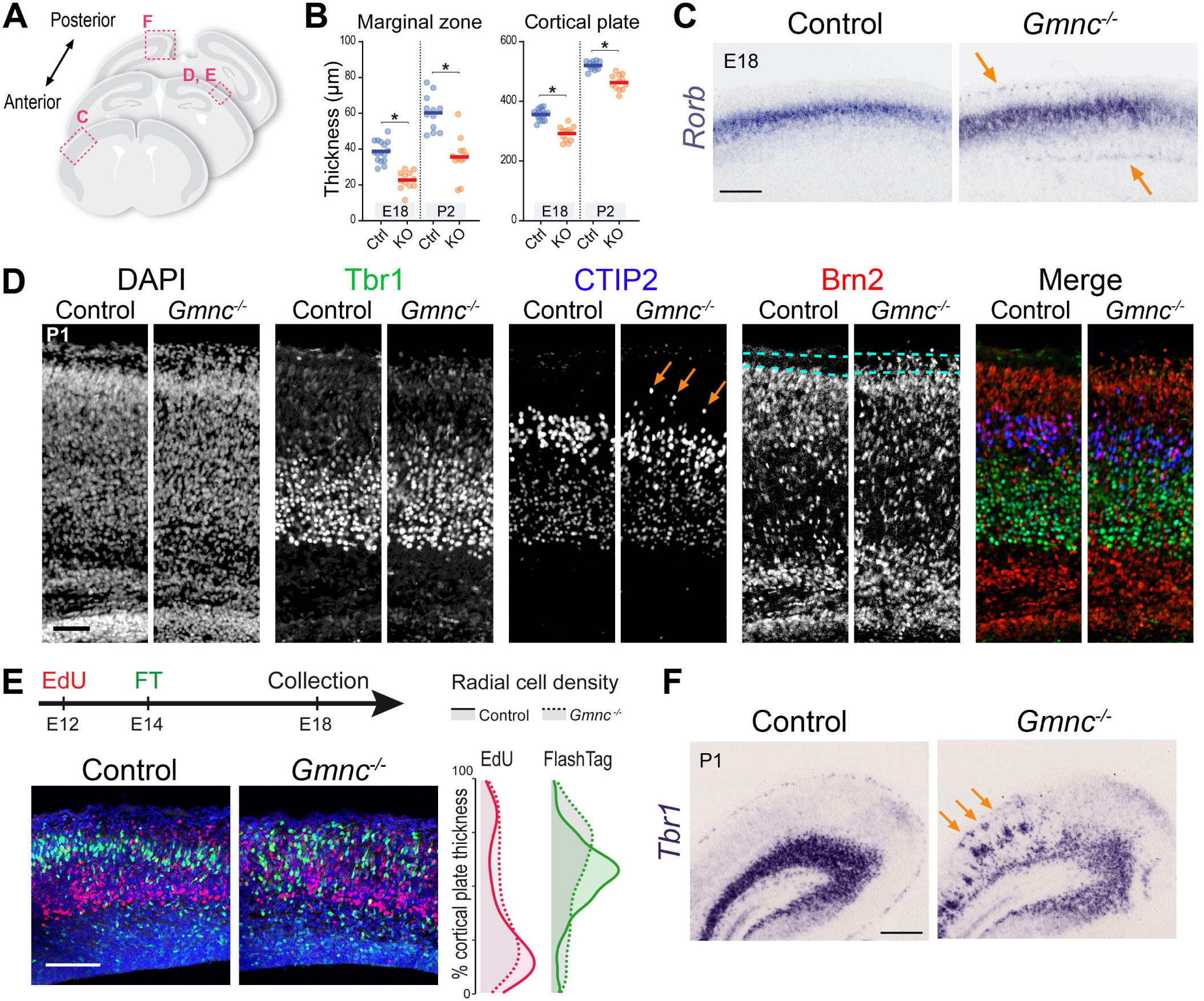
Radial migration defects upon CRs depletion. (A) Drawing indicating the regions shown in B-G. (B) Quantification of the cortical plate and MZ thickness in control and *Gmnc* mutants at E18 and P2 (3 animals and 4 sections per animal were measured for each stage and genotype). * p<0.0001 using Mann Whitney test. (C) *In situ* hybridization for *Rorb* in a rostral section of the cortex from E18 control and *Gmnc^-/-^* embryos. Arrows point to ectopic cells in the MZ and subplate. (D) Immunostaining for Tbr1, CTIP2 and Brn2 in a cortical column of the presumptive somatosensory area from P1 control and *Gmnc^-/-^* mice. Arrows point to ectopic CTIP2^+^ cells, dashed lines delineate the MZ that is invaded by Brn2^+^ cells in mutants. (E) Cortical column of the presumptive somatosensory area from E18 control and *Gmnc^-/-^* mice following labelling of E12- and E14-born neurons with EdU and FlashTag, respectively. Quantification (right) showing the density of labelled cells along the cortical plate thickness. Dashed lines correspond to mutants. >3200 cells considered, no less than 680 per condition from 3 animals per genotype. (F) *In situ* hybridization for *Tbr1* in the dorsal and caudal cortex (presumptive visual area) of P1 control and *Gmnc^-/-^* mice. Arrows point to clumps of ectopically positioned cells in mutants. Scale bars: 200µm in C, F, 50µm in D, 100µm in E.

**Figure 4.**
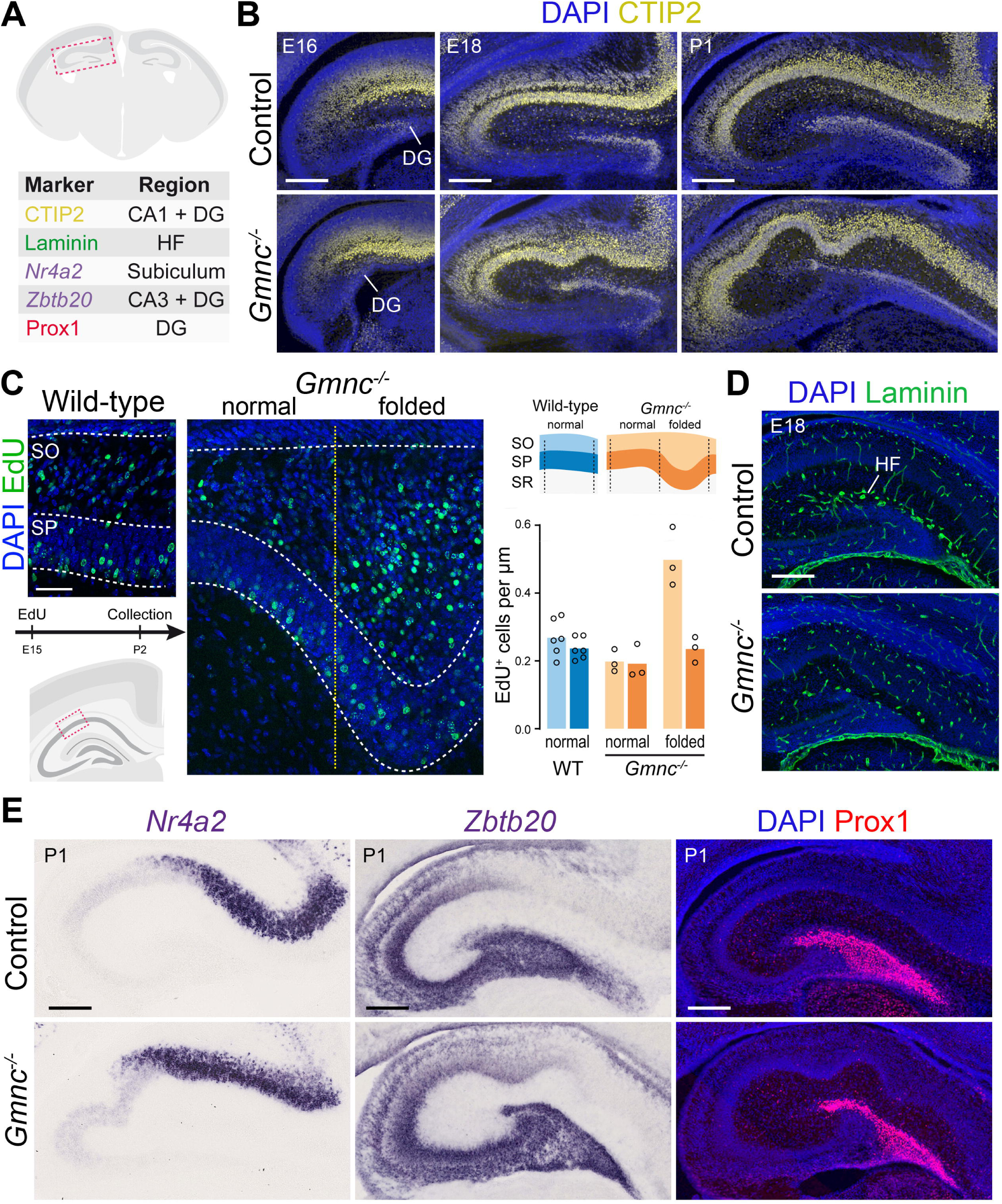
Hippocampal morphogenesis defects upon CRs depletion. (A) Drawing indicating the region shown in B-E (top) and region specificity of the markers used (bottom). (B) Immunostaining for CTIP2 on coronal sections of the hippocampus from E16 to P1 in control and *Gmnc^-/-^* animals. DG: dentate gyrus anlage. (C) Immunostaining for EdU on sections of the hippocampus at P2 in control and *Gmnc* mutants following EdU incorporation at E15. SO: *stratum oriens*, SP: *stratum pyramidale*. The yellow dashed line separates the regions considered normal and folded in mutants. Quantification of EdU labelled cells in control and mutant (n=3 each), considering normal or folded regions of CA1. (D) Immunostaining for Laminin (green) on coronal sections of the hippocampus in E18. HF: hippocampal fissure, only distinguishable in controls. (E) *In situ* hybridization for *Nr4a2* and *Zbtb20*, and immunostaining for Prox1 on coronal sections of the hippocampus at P1 in control and *Gmnc^-/-^* animals. Scale bars: 200µm in B, C, E. 50µm in D.

## Results

### Spatiotemporal dynamics of CRs depletion in *Gmnc* mutants

We have previously shown that upon loss of the multiciliogenesis master gene *Gmnc*, septum-, ThE- and hem-derived CRs are produced in apparently normal numbers, migrate correctly to cover the dorsal cortex, but fail to express their usual marker *Trp73*, initiate apoptosis from E12 onwards and are completely absent at E14 (Moreau et al., 2023). By contrast, VP-derived CRs never express *Gmnc* and are expected to remain unaffected by its loss. We further characterized the spatiotemporal dynamics of CRs depletion in *Gmnc* mutants. To this end, we performed *in situ* hybridization for *Nhlh2*, a transcription factor expressed by all CRs subtypes at E12 (Fig 1A) and highly specific of CRs from E13 onwards (Fig. 1B).

At E14, the dorsal cortex MZ of *Gmnc* mutants was completely devoid of *Nhlh2*^+^ cells, whereas the lateral cortex showed normal staining (Fig. 1C). This phenotype was already observed at E13, although residual staining could still be observed in the dorsal cortex, but not at E12 (Fig S1A). At E16 and E18, only very few *Nhlh2*^+^ cells were detected in the lateral cortex of mutants (Fig 1D and S1B). To rule out the possibility that lateral CRs simply loose the expression of *Nhlh2* with time, we performed *in situ* hybridization for three additional pan-CRs genes, namely *Tbr1*, *Lhx5* and *Calb2* (Fig 1A). In all cases, we could only find a few positive cells scattered in the lateral cortex MZ of mutants (Fig 1E), indicating lateral CRs progressively disappear.

In the hippocampal region of E18 *Gmnc* mutants, we observed *Nhlh2*^+^ cells along the hippocampal fissure and pial surface of the dentate gyrus (Fig. 1F), indicative of CRs persistence. We and others previously showed that p73 expression in the brain is lost upon *Gmnc* inactivation (Lalioti et al., 2019; Moreau et al., 2023). However, we performed *in situ* hybridization for *Trp73* in mutants, and observed positive cells at E18 that were not present at E14 (Fig. 1F, G). These cells also expressed P73 protein and their density was reduced ∼5 fold in mutants compared to controls (56 ± 13 and 260 ± 32 cells per mm of HF, respectively, Fig. 1H, I). They were found Tomato^+^ upon genetic tracing with *Wnt3a^Cre^*, suggesting a *bona fide* hem-derived CRs identity (Fig. 1J). Because p73 expression is completely absent at E13 and E14 in *Gmnc^-/-^* embryos (Fig 1G and (Moreau et al., 2023)), we concluded that the few hippocampal CRs observed in E18 mutants most likely correspond to a cohort generated at late stages in a *Gmnc*-independent manner. These cells were confined to the hippocampus and did not migrate in the dorsal cortex as indicated by genetic tracing (Fig. 1K).

Taken together, these data show that in *Gmnc* mutants the dorsal cortex is completely devoid of CRs from E14 onwards, the lateral cortex shows normal CRs density at E14 but later gets depleted progressively, whereas in the hippocampal formation, a reduced contingent of late-born CRs remains present (Fig 1L).

### Depletion of dorsal cortex CRs leads to a dramatic loss of Reelin expression

We then evaluated the extent to which Reln protein distribution is affected by CRs loss in *Gmnc* mutants. At E14, consistent with the extent of CRs depletion (Fig 1C), normal Reln immunoreactivity was observed in the lateral cortex, whereas only very few Reln^+^ cells were detected in the dorsal cortex (Fig 2A). We have previously shown that these remainders correspond to mis-specified and often ectopically positioned (deep in the cortical plate) hem-derived CRs undergoing apoptosis (Moreau et al., 2023). Reln^+^ cells of the lateral MZ were found Foxg1-negative, confirming their CRs identity (Fig 2B). Despite the progressive disappearance of lateral CRs, we found Reln protein levels to be maintained at E16 and E18 in the lateral cortex MZ of mutants, leading to the formation of a sharp boundary between lateral and dorsal MZ (Fig 2C). At perinatal stages, the sharp distinction between the massive decrease in Reln immunoreactivity of the dorsal cortex MZ of mutants, and the almost normal immunoreactivity of the lateral cortex was even more striking (Fig 2C, D, E and S2). Reln levels appeared maintained in the lateral cortex MZ, but also by superficial neurons of the insular and piriform cortex, especially at caudal levels (Fig 2D, S2 also illustrated in 5G). In addition, sparse Reln^+^ cells could be detected in the cortical plate of both controls and mutants at these stage, reminiscent of cortical interneurons. Finally, Reln immunoreactivity in the hippocampus of *Gmnc* mutants at E18 was found strongly reduced compared to controls, but not completely abolished, consistent with the presence of late-born CRs (Fig 2F).

Taken together, these data show that the amounts and distribution pattern of Reln protein are severely affected in *Gmnc* mutants from midcorticogenesis onwards, with the dorsal cortex displaying a more complete loss than the hippocampus and the lateral cortex appearing close to normal (Fig 2G).

### Defects in radial migration arrest upon depletion of dorsal cortex P73^+^ CRs

We first decided to investigate the consequence of P73^+^ CRs depletion in the dorsal cortex. Since *Gmnc* mutants develop hydrocephalus leading to increased mortality during the first postnatal weeks (Terré et al., 2016), we conducted most of our analysis at perinatal stages, between E18 and P2. At such stages, we failed to detect any ventricular enlargement in mutants compared to controls (Figs. 2D, S2). Using DAPI staining, we found the MZ and cortical plate significantly thinner (∼40% and ∼15% decrease, respectively) in mutants compared to wild-type littermates (Fig. 3B), indicating that the severe reduction of cortical thickness observed in adults (Terré et al., 2016) is not only secondary to hydrocephalus. Within the cortical plate, only layers 2-4 were found significantly thinner in mutants compared to controls (Fig S3A). Upon staining for layer-specific markers, we observed a number of anomalies in mutants. At rostral levels of the somatosensory region, the presumptive layer 4, labelled by *in situ* hybridization for *Rorb*, appeared less compact, and ectopically positioned cells were found in the subplate and MZ (Fig. 3C). In the presumptive somatosensory area, immunostaining for Tbr1 (layer 6), CTIP2 (layer 5) and Brn2 (layers 2-4) indicated a global preservation of the radial lamination pattern in mutants, although the delineation of each layer appeared less sharp, as exemplified by the presence of ectopic CTIP2^+^ cells in upper layers (Fig 3D). A robust defect observed in mutants was the invasion of the MZ by Brn2^+^ cells (Fig 3D, also evidenced with Foxg1 staining Fig S3B), suggesting a failure to stop radial migration. Since these defects are suggestive of an impairment in radial migration in *Gmnc^-/-^*embryos, we performed EdU birthdating at E12 followed by intraventricular FlashTag injection at E14, in order to successively label early-born and late-born neurons, respectively. Analysis at E18 indicated that the two cohorts were well segregated in wild-type brains whereas they appeared more intermingled in mutants (Fig. 3E). In addition, quantifications indicated that FlashTag^+^ E14-born cells reach more superficial positions in mutants than controls, confirming a defective termination of the radial migration process. Thus, despite the severe depletion of CRs and Reln in *Gmnc* mutants, E14-born neurons retained their ability to migrate past earlier born neurons and rather failed to stop. By contrast, the radial positioning of Prox1^+^ GABAergic cortical interneurons was found unaffected upon P73^+^ CRs depletion (Fig S3C). Finally, in the presumptive visual area of mutants, we observed clumps of *Tbr1*-expressing cells ectopically positioned throughout the radial dimension instead of being confined to deep layers (Fig. 3F), indicating that not all cortical regions are equally sensitive to CRs and Reln loss. Overall, our data indicate that premature depletion of P73^+^ CRs from the dorsal cortex at midcorticogenesis results in alterations of radial migration of glutamatergic neurons that differentially affect distinct areas.

### Hippocampal morphogenesis defects upon CRs depletion

Adult *Gmnc* mutants were previously shown to harbour marked hippocampal dysgenesis (Terré et al., 2016) but it remained unclear whether the defects resulted from the severe hydrocephalus or not. In addition, mouse mutants for *Trp73*, one of the major downstream effector of Gmnc, were previously shown to lack a HF (Meyer et al., 2004; Meyer et al., 2019). We therefore evaluated the impact of CRs premature loss in *Gmnc* mutants on hippocampal morphogenesis (Fig. 4A). At E16, CTIP2 immunostaining allowed us to distinguish between the dentate gyrus (DG) anlage and presumptive CA1 neurons, as the HF starts to separate the two regions. In *Gmnc* mutants, such a distinction proved difficult to make (Fig. 4B). At E18 and P1, we observed the progressive appearance and worsening of an abnormal folding of the prospective CA1 region in mutants (Fig. 4B). We reasoned this could be due to excessive proliferation and therefore performed EdU pulse labelling at E15, to find a ∼2-fold increase in the density of EdU^+^ cells in the *stratum oriens* (but not *stratum pyramidale*) of folded regions compared to wild-type or unfolded regions in mutants (Fig 4C). These data suggest that CRs are able to regulate the proliferation of hippocampal progenitors. Furthermore, we performed Laminin immunostaining to visualize the basal lamina of the HF and endothelial cells. We observed a clear decrease in mutants, confirming that CRs are required for HF formation but also pointing to an unexpected role for CRs in vasculature ingrowth (Fig 4D). Finally, we performed *in situ* hybridization for *Nr4a2*, to label the subiculum, *Zbtb20*, showing high expression in CA3 and DG, and immunostaining for Prox1 to visualize the DG, and observed an apparently normal parcellation of hippocampal territories and specification of neuronal types (Fig. 4E). Overall, our data confirm that hippocampal CRs are required for HF formation, point to an additional role in the control of progenitor proliferation and vasculature growth, and suggest CRs are dispensable for patterning of the hippocampal formation.

### Defects in the lateral cortex upon P73^+^ CRs depletion

We noticed an unusual cellular organisation in the lateral aspect of *Gmnc* mutant cortices that was most prominent at late gestational stages and intermediate antero-posterior levels (Fig. 5A). At the prospective boundary between the insular and dorsal cortex, the Reln-immunoreactive and cell-body poor MZ of the lateral cortex appeared to dive within the tissue in mutants (Fig 5B). Foxg1 immunostaining illustrated how cortical neurons appear excluded from the ingrowing Reln-rich region in mutants (Fig 5C). This was paralleled by the ectopic positioning of Nrp1^+^ fibres, presumably olfactory axons that normally extend over the surface of the lateral cortex, and were found to form a fissure in mutants (Fig. 5D). Using *in situ* hybridization for *Rorb* and *Nr4a2* to label deep insular neurons and claustrum, respectively, we observed a clear displacement suggesting cells were pushed deeper than normal within the depth of the lateral cortical plate (Fig. 5E-G). *In situ* hybridization for *Reln* also allowed to visualize the abnormal bending of superficial layers of the insular cortex in mutants (Fig 5G). Using Ctip2 and Tbr1 immunostaining to further visualize the arrangement of neuronal cell types in the affected region of mutants, we were able to delineate insular and dorsal cortex layered organization (Fig 5H-I). We found the global structure preserved but distorted and displaced in mutants, in a manner suggesting physical pressure or repulsion from the Reln-rich region. Alternatively, the observed tissue reorganization could stem from the overmigration of dorsal cortex neurons, resulting in the apparent engulfment of the lateral cortex MZ. Nevertheless, we concluded that the discontinuity in Reln distribution between the lateral and dorsal MZ of mutants disrupts local cell positioning and results in apparent tissue bending.

**Figure 5.**
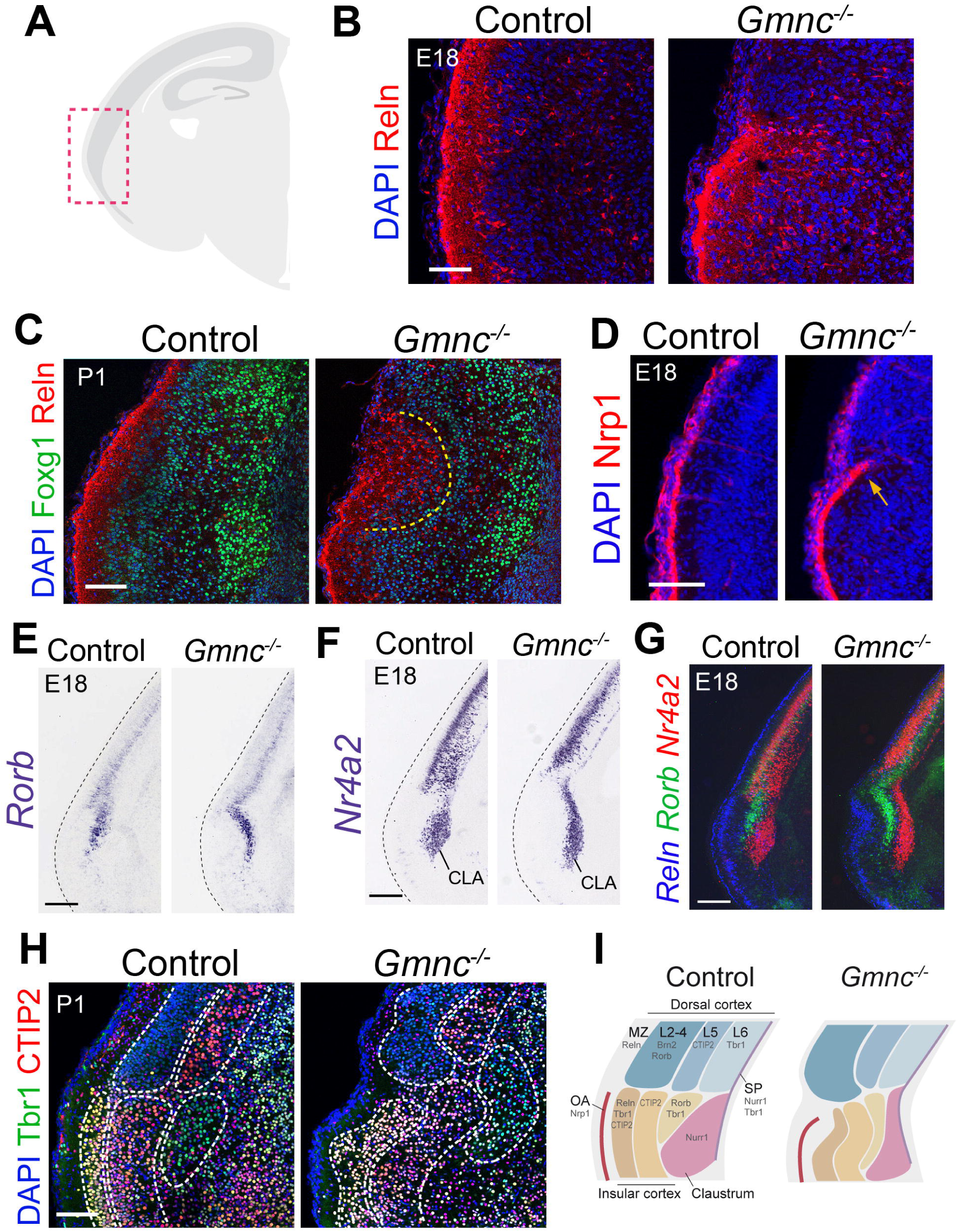
CRs depletion results in defects in the lateral cortex. (A) Drawing indicating the region shown in B-H. (B) Immunostaining for Reln in the lateral cortex at E18 in control and *Gmnc^-/-^* embryos illustrating the sharp boundary between insular and dorsal cortex. (C) Immunostaining for Reln and Foxg1 in the lateral cortex of P1 control and *Gmnc* mutants. The dashed line surrounds the Reln-rich and DAPI-poor region. (D) Immunostaining for Nrp1 in the lateral cortex of E18 control and *Gmnc^-/-^*embryos. The arrow points to presumptive olfactory fibres ectopically invading the cortical parenchyma in mutants. (E-F) *In situ* hybridization for *Rorb* (E) and *Nr4a2* (F) in the lateral cortex of E18 control and *Gmnc^-/-^* mice. CLA: claustrum. (G) Overlay of three adjacent serial sections from the same embryo processed for *Reln*, *Rorb* or *Nr4a2 in situ* hybridisation. (H) Immunostaining for CTIP2 and Tbr1 at P1 in control and *Gmnc^-/-^* animals. The dashed lines delineate layers of the dorsal and insular cortex. (I) Illustration of the defects observed. OA: olfactory axons, SP: subplate. Scale bars: 100µm in B-D, H. 200µm in E-G.

### Cell type composition is preserved upon P73^+^ CRs depletion

To better appreciate whether cell specification might be affected by CRs loss, we implemented single-cell RNAseq experiments. We dissected the entire cortex, including the lateral cortex and the hippocampal formation, from four wild-type and four *Gmnc^-/-^* P0 newborn animals and generated one library for each genotype (Fig. 6A). After quality control, cell filtering and integration, we obtained a dataset containing 24,526 cells, 29.8% originating from wild-type embryos, and a median 2,797 genes detected per cell. Clustering and dimensionality reduction allowed to identify cell classes on the basis of well-known marker genes expression (Fig. 6B, C). Contribution of wild-type and mutant cells was fairly constant across cell classes, indicating no major composition bias (Fig. 6D). However, displaying the genotype of each cell on the 2D embedding revealed two small clusters, almost exclusively composed of wild-type cells (Fig. 6E). The first one was found among excitatory neurons and showed high *Trp73* and *Reln* expression, reminiscent of CRs identity. The second was found among glial cell types and showed *Trp73* and *Ccdc67* (also known as *Deup1*), indicating a multiciliated ependymal cell identity. To further confirm these findings, we sub-clustered excitatory neurons and glial cells separately and performed dimensionality reduction (Fig. 6F, J). Marker gene expression allowed the identification of major neuronal excitatory types (Fig. 6G), including CRs that form a well-defined and isolated cluster. 94.1% of CRs present in the dataset (32 out of 34 cells) originated from the wild-type condition (Fig. 6H, I), far above other clusters that ranged around 30%. We therefore confirmed that CRs are almost, but not completely absent in *Gmnc* mutants at birth. Regarding glial cells, we identified a cluster of ependymal cells that diverged from all other clusters (Fig. 6J, K), and was only composed of wild-type cells (Fig. 6J-M). Thus, ependymal cells not only fail to grow multiple cilia in *Gmnc* mutants, but also lose a large part of their transcriptomic identity. Overall, the only clusters showing overabundance of mutant cells were made of doublets that escaped cell filtering (illustrations embedded in our codes at https://fcauseret.github.io/P0_GmncKO/) and probably resulted from the larger number of cells encapsulated in the mutant condition compared to control. We therefore concluded that the morphological defects observed in *Gmnc^-/-^* animals rather result from the abnormal organisation of otherwise normal cell types than from the specification of aberrant cell identities.

**Figure 6.**
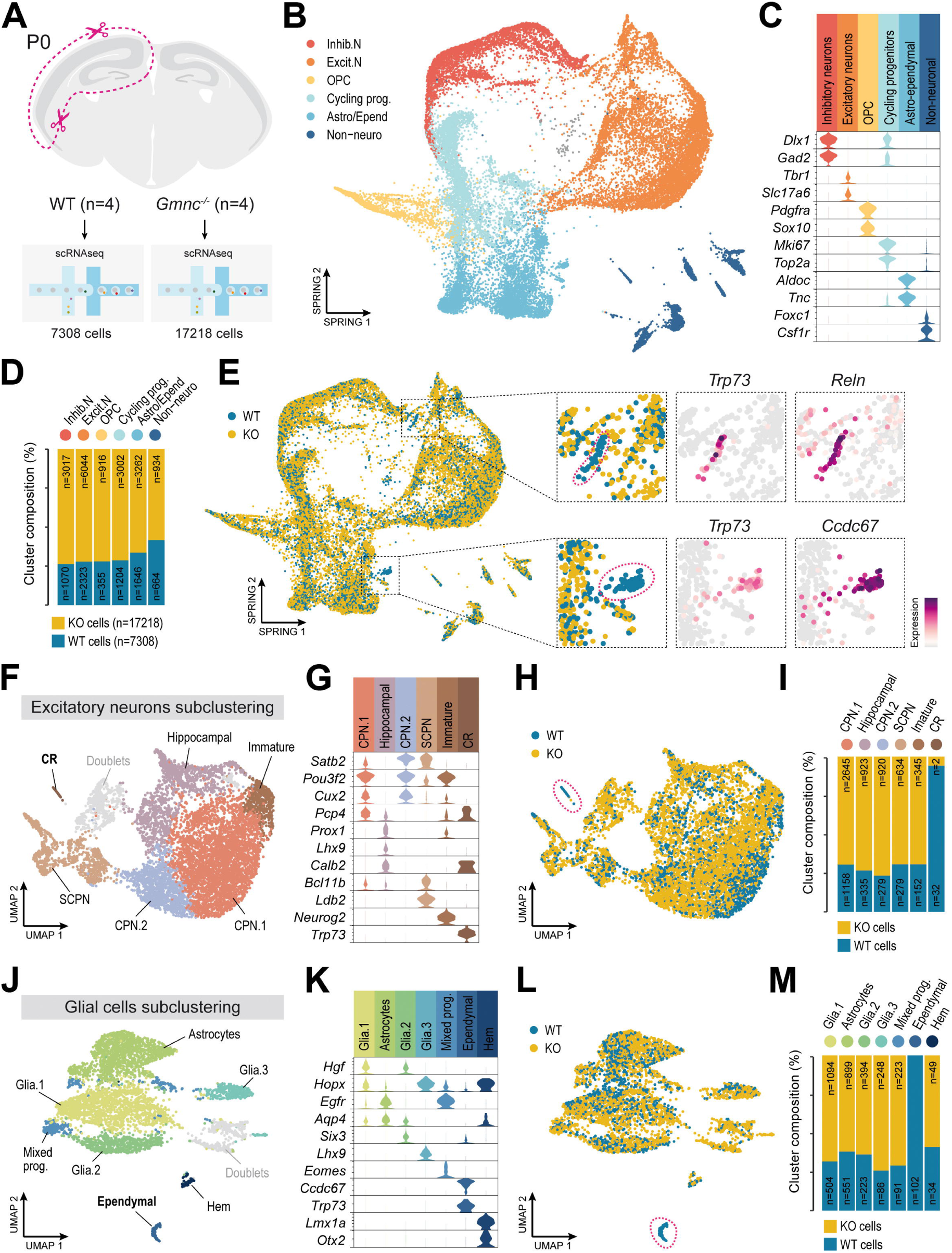
Cell type composition in Gmnc mutants. (A) Experimental design of the single-cell RNAseq approach. (B) 2D representation of the data following SPRING dimensionality reduction. Cells (points) are color-coded according to identity classes. (C) Violin plots of selected markers used for cell-type annotation in B. (D) Stacked histogram depicting the relative contribution of wild-type and *Gmnc^-/-^* cells to each cell class. The number of cells is indicated on each bar. (E) Dimensionality reduction plot showing cells color-coded according to their genotype. The magnified boxes indicate two small clusters (dashed lines) lacking *Gmnc^-/-^* cells. The expression of selective marker genes for these two clusters is shown. (F) Dimensionality reduction plot following excitatory neurons subclustering. (G) Violin plots of selected markers used for cell-type annotation in F. (H) Dimensionality reduction plot showing excitatory neurons color-coded according to their genotype. The dashed line surrounds the cluster of CRs that is almost exclusively composed of wild-type cells. (I) Stacked histogram depicting the relative contribution of wild-type and *Gmnc^-/-^* cells to each neuronal type. Note the strong composition bias among CRs relative to other clusters. (J) Dimensionality reduction plot following glial cells subclustering. (K) Violin plots of selected markers used for cell-type annotation in J. (L) Dimensionality reduction plot showing glial cell types color-coded according to their genotype. The dashed line surrounds the cluster of ependymal cells that is exclusively composed of wild-type cells. (M) Stacked histogram depicting the relative contribution of wild-type and *Gmnc^-/-^* cells to each glial type. Note the outstanding composition of ependymal cells relative to other clusters.

### Reln depletion from P73^+^ CRs only partially recapitulate CRs loss

CRs are tightly associated with Reln expression but so far, no direct comparison of the consequences of CRs loss vs Reln loss has been reported. We therefore attempted to conditionally inactivate *Reln* in P73^+^ CRs and determine whether it would recapitulate some of the defects observed in *Gmnc* mutants. We used the *ΔNp73^Cre^* line that target CRs originating from the hem, septum and thalamic eminence (Tissir et al., 2009) in either *Reln^lox/+^*(controls) or *Reln^lox/-^* (cKO) littermates. All animals additionally carried a *Rosa26^tdTomato^* allele in order to monitor recombined CRs and distinguish them from other Reln^+^ cells, mostly GABAergic interneurons that have invaded the MZ at late gestational stages (Fig. 7A, B, S4B). In some instances, we also used a full deletion of *Reln*, equivalent to the *Reeler* mutant (Falconer, 1951), using the *PGK^Cre^* strain (Lallemand et al., 1998).

**Figure 7.**
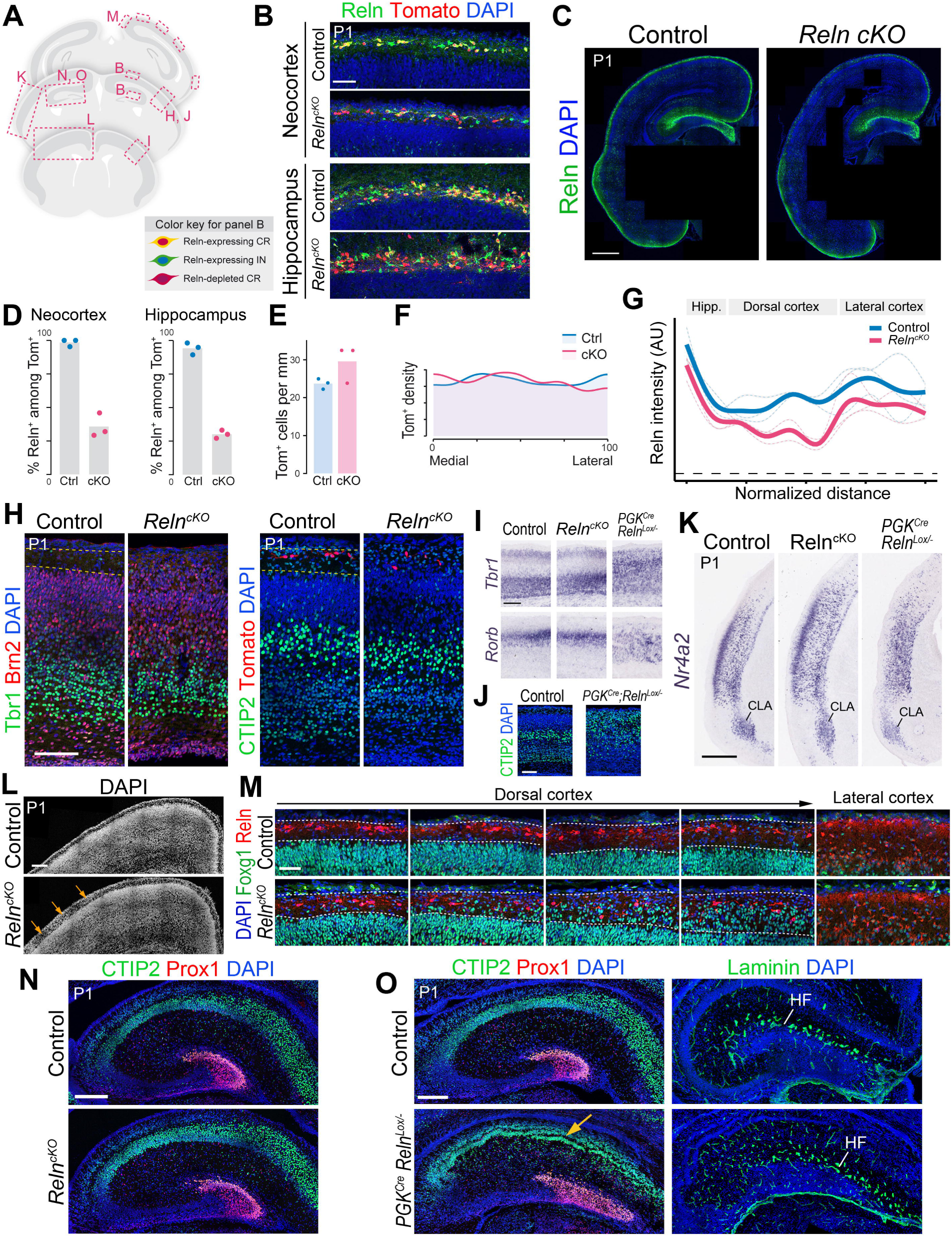
CRs-specific *Reln* deletion affects radial migration. (A) Drawing indicating the regions shown in B-L (top) and color key for the panel shown in B. (B) Immunostaining for Reln and Tomato in the neocortex or hippocampus of P1 control or *Reln^cKO^*animals. (C) Immunostaining for Reln in control and cKO showing the decreased Reln expression, in the dorsolateral cortex and hippocampus. (D) Quantification of the fraction of Reln^+^ cells among Tomato^+^ CRs in the dorsal cortex and hippocampus. Each point corresponds to one animal. (E) Quantification of the density of Tomato^+^ CRs in the dorsal cortex MZ of control and cKO mice. Each point corresponds to one animal. (F) Density of Tomato^+^ CRs along the medio-lateral axis of the cortical MZ (n=3 animals per genotype). (G) Quantification of the fluorescence intensity of Reln (arbitrary units) along the HF and cortical MZ. The dashed lines represent each animal considered (n=3 per genotype). (H) Immunostaining for Tbr1and Brn2, or for CTIP2 and Tomato in a cortical column of the presumptive somatosensory cortex of P1 control and *Reln^cKO^*. (I) *In situ* hybridization for *Tbr1* and *Rorb* in a rostral section of the cortex from P1 control, *Reln^cKO^* and *PGK^Cre^;Reln^lox/-^* mice. (J) Immunostaining for CTIP2 in a cortical column of the presumptive somatosensory cortex of P1 control and *PGK^Cre^;Reln^lox/-^* mice. (K) *In situ* hybridization for *Nr4a2* in the lateral cortex of P1 control, *Reln^cKO^* and *PGK^Cre^;Reln^lox/-^* mice. CLA: claustrum. (L) DAPI staining of nuclei in the rostromedial cortex of P1 control, *Reln^cKO^* and *PGK^Cre^;Reln^lox/-^* mice. Arrows point to cells abnormally positioned in the MZ of mutants. (M) Immunostaining for Foxg1 and Tomato along the medio-lateral axis of the caudal cortex of P1 control or *Reln^cKO^*. (N) Immunostaining for CTIP2 and Prox1 in the P1 hippocampus of control and *Reln^cKO^*. (O) Immunostaining for CTIP2 and Prox1 or Laminin in the P1 hippocampus of control and *PGK^Cre^;Reln^lox/-^* mice. The arrow points to the defect in compaction of the CA1 pyramidal layer. HF: hippocampal fissure. Scale bars: 50µm in B, K. 100µm in F, H. 200µm in G, J, L, M. 500µm in I, K.

In cKO, we found Reln immunostaining in the MZ decreased compared to controls (Fig 7B, C) indicating efficient recombination. However, only two third of Tomato^+^ targeted CRs were found confidently negative for Reln immunoreactivity (Fig 7B, D, S4B). This would imply a partially ineffective recombination at the *Reln* locus relative to the *Rosa26* locus. Nevertheless, quantification of Reln staining intensity throughout the HF and cortical MZ indicated a reduction in cKO that was more pronounced in the dorsal cortex compared to other regions (Fig 7G), although it did not match the extent of the almost complete loss observed in *Gmnc* mutants (compare with Fig 7C and 2B, or 7G and 2C). By contrast, both the number and distribution pattern of Tomato^+^ CRs along the medio-lateral axis were found unchanged between genotypes (Fig 7E, F, S4A) confirming that Reln itself does not influence CRs production, migration, distribution or survival (Anstötz et al., 2019). We therefore concluded that cKO display a significant but incomplete Reln loss in P73^+^ CRs without visible alteration of CRs numbers or position.

We first investigated neocortical radial organisation in *Reln^cKO^*. We could not clearly distinguish cKO from controls using markers of layer 4-6 *Rorb*, CTIP2 or Tbr1 (Fig 7H-J), contrasting with the major disorganization observed in full *PGK^Cre^* ; *Reln^lox/-^* mutants (Fig 7I, J), indicating that reduced amounts of Reln are sufficient to ensure early-born neurons positioning. However, immunostaining for the superficial layer marker Brn2 allowed to visualize neurons invading the MZ in cKO (Fig 7H), highly reminiscent of *Gmnc* mutants (Fig 3D). This phenotype was found less prominent in the medial cortex (Fig. 7L), correlating with the amounts of Reln in the MZ (Fig 7C, G), and was especially highlighted using Foxg1 immunostaining, labelling all telencephalic neurons except CRs (Fig 7M). Importantly, overshooting neurons were often observed in close proximity of either CRs or Reln^+^ cells (Fig 7M and S4C), suggesting they were sensitive to the total amount of Reln present in the MZ rather than some sort of diffusion gradient formed around secreting cells. Consistently, we did not observe MZ invasion in the lateral cortex of cKO, where Reln^+^/Foxg1^+^ superficial insular neurons were normally positioned (Fig 7M and S4D). Thus, conditional Reln deletion failed to recapitulate P73^+^ CRs loss in the lateral cortex, as further evidenced by the normal shape and position of the claustrum shown by *in situ* hybridization for *Nr4a2*, whereas in full *PGK^Cre^* ; *Reln^lox/-^* mutants the claustrum occupied an aberrant superficial position (Fig 7K). One possible interpretation of these data is that the remaining Reln^+^ cells present in the dorsal cortex MZ of cKO prevent the formation of a sharp boundary at the interface between lateral and dorsal cortex. However, we cannot rule out the alternative that P73^+^ CRs regulate lateral cortex morphogenesis independently of Reln. Finally, despite a measureable decrease in Reln immunoreactivity in the hippocampus (Fig 7B, S4E), Reln cKO were undistinguishable from controls following CTIP2 and Prox1 immunostaining (Fig 7N). Since the magnitude of Reln inactivation in hippocampal CRs of the cKO is similar to that of CRs depletion in the hippocampus of *Gmnc* mutants (compare Figs 2E and 7D), we concluded that CRs implication in hippocampal morphogenesis is, at least in part, Reln-independent. This is also supported by the observation that full *PGK^Cre^* ; *Reln^lox/-^* mutants, although failing to form a compact pyramidal layer as previously described (Hamburgh, 1963), never display abnormal CA1 folding as reported for *Gmnc* mutants, and develop a HF that is normally invaded by vasculature, according to Laminin staining (Fig 7O, compare to Fig 4C).

Taken together, these results point to a function at mid-corticogenesis for P73^+^ CRs-derived Reln in arresting the radial migration of upper layers’ neurons, in addition to Reln-independent functions for hippocampal CRs in the control of progenitor proliferation and vasculature growth.

## Discussion

For a century following their initial description by Santiago Ramón y Cajal and Gustaf Retzius, CRs remained a poorly studied cell type. They became the focus of attention only from the discovery of Reln in 1995 (D’Arcangelo et al., 1995; Ogawa et al., 1995). Since then, CRs became (and remain) tightly associated to Reln. CRs express high levels of Reln, and their strategic location at the surface of the developing cortex from early stages of corticogenesis led to the proposal that CRs-derived Reln controls radial migration (Ogawa et al., 1995). However, this textbook model is not entirely satisfactory. Indeed, other sources of Reln exist, including GABAergic cortical interneurons, glutamatergic neurons of the insular and piriform cortex and immature migrating glutamatergic neurons in the neocortex. Conditional deletion of *Reln* from CRs or cortical interneurons recently demonstrated that both sources actually cooperate to fine-tune the inside-out arrangement of neocortical layers (Vílchez-Acosta et al., 2022). To complement these findings, we took advantage of the *Gmnc* mutant model that targets most CRs subtypes (P73^+^ originating from the hem, septum and ThE), to address the consequence of their removal at midcorticogenesis and compare to Reln loss.

### CRs depletion and Reln deletion

We have previously shown that *Gmnc* mutants phenocopy *Trp73* mutants, regarding CRs differentiation at least (Meyer et al., 2004; Moreau et al., 2023), consistent with p73 being the main downstream effector of Gmnc (Lalioti et al., 2019). In addition, *Trp73* mutants display a thinner dorsal cortex, abnormal hippocampal folding, absence of HF, as well as macroscopic defects around the rhinal fissure (Meyer et al., 2004), all features highly reminiscent of those we described in *Gmnc* mutants. Since none of these mutants recapitulate the severe layer disorganization observed in *Reeler* mutants (Boyle et al., 2011), one could argue that a minimal number of CRs or CRs-derived Reln are sufficient to ensure neocortical lamination. However, we have previously shown that the initial production and migration of CRs occurs normally in *Gmnc* mutants (Moreau et al., 2023), allowing CRs presence in the MZ up to E12. Although misspecified, these cells retain Reln expression, likely favouring early processes such as preplate splitting and opening the possibility that the early presence of CRs and Reln in *Gmnc* mutants contribute to lamination at later stages. Consistent with this, our results point to the importance of CRs-derived Reln to ensure the correct arrest of migration of later born neurons, resulting in their invasion of the MZ upon depletion of CRs in *Gmnc* mutants or of Reln in cKOs. Preplate splitting occurs in *Emx1^Cre^;Wnt3a^DTA^* embryos (Yoshida et al., 2006) despite an almost complete ablation of the hem prior to the temporal window of CRs production. In addition, these mutants display a fairly normal lamination, at least in rostral regions where septum-derived CRs are present. Although overmigration was not directly assessed in these mutants, the authors proposed that a majority of CRs are dispensable for neocortical lamination, and argue for a compensatory effect of Reln production by early post-mitotic glutamatergic neurons (Yoshida et al., 2006). A model that is currently well supported by experimental data would be that distinct sources of Reln (CRs, early postmitotic glutamatergic cortical neurons and GABAergic cortical interneurons) cooperate in space and time to regulate the inside-out radial migration (Vílchez-Acosta et al., 2022; Yoshida et al., 2006). Yet, neither the half-life of Reln, nor its ability to diffuse *in vivo* are precisely known, further blurring our understanding of the quantity and distribution that are precisely required to ensure proper corticogenesis. Furthermore, the caudal cortex of hem-ablated embryos was found highly disorganized, in a manner reminiscent of what we observed in *Gmnc* mutants (compare Fig. S1 of (Yoshida et al., 2006) with our Fig. 3F), suggesting a high degree of cell-type and/or area specificity when considering the impact of CRs to cortical development. This is also the case for Reln as the magnitude of lamination defects in the *Reeler* mutants are variable across brain regions (Boyle et al., 2011), and the response to ectopic Reln overexpression by *in utero* electroporation differs along the anteroposterior axis (Riva et al., 2024).

### Reln-independent functions of CRs

The idea that CRs could provide the developing brain with secreted signals distinct from Reln initially stemmed from *in vitro* graft experiments showing that both wild-type and *Reeler*-derived CRs could equally regulate the phenotype of radial glia in cerebellar slices (Soriano et al., 1997). Additional Reln-independent functions for CRs were also proposed in establishing hippocampal connectivity (Del Río et al., 1997), or controling neocortical progenitor proliferation and cortical arealization by CRs subtypes distribution (Griveau et al., 2010). However, the demonstration that P73^+^ CRs depletion in *Gmnc* mutants affects hippocampal morphogenesis more severely than CRs-specific or even complete *Reln* inactivation provides a direct evidence supporting a Reln-independent function for CRs. Our work also corroborates previous observations that *Trp73* mutants lack a HF, unlike conditional or constitutive *Reln* mutants (Anstötz et al., 2019; Meyer et al., 2004; Vílchez-Acosta et al., 2022). Transcriptomic analyses indicated CRs secrete a range of signalling molecules beyond Reln, that are likely to impact cortical development (Causeret et al., 2021; Griveau et al., 2010; Yamazaki et al., 2004). Future studies will be required to assess to which extent such candidates contribute to Reln-independent functions for CRs.

### Role of CRs in tissue morphogenesis

One speculation that could be put forward in the light of our findings is the possibility that CRs control tissue bending as observed in the mammalian hippocampus. Contrary to mammals, sauropsid species (birds, turtles, lizards) show little, if any, CRs at the surface of their medial pallium (Bar et al., 2000; Cabrera-Socorro et al., 2007; Moreau et al., 2023; Tissir et al., 2003) and lack a HF (Hevner, 2016). Yet, cell-type conservation studies indicate that homologs to CA1, CA3 and DG neurons can be found across amniotes (Tosches et al., 2018; Zaremba et al., 2024). Furthermore, in mouse and humans hem-derived CRs accumulation occurs precisely where the HF will later form (Hevner, 2016; Meyer et al., 2004; Meyer et al., 2019). It is therefore tempting to speculate that the increased CRs density in the medial pallium of mammals contributed to the evolutionary emergence of the HF and subsequent folding of the hippocampal formation. In this line, it is worth mentioning that a subset of CRs was found to localize preferentially at the bottom of sulci in the human brain, suggesting a function in cortical folding (Meyer and González-Gómez, 2018a; Meyer and González-Gómez, 2018b). In addition, our data on the lateral cortex suggest that discontinuity in Reln distribution could also be a way to achieve tissue bending. Because the biological activity of RELN variants found in patients with cortical malformations can be correlated with the gyration pattern (pachygiria for dominant-negative variants vs polymicrogyria for gain-of-function variants) (Riva et al., 2024), both CRs and Reln appear as potential players in cortical folding.

### Unmasking VP-derived CRs

VP-derived CRs lack a specific positive transcriptomic signature and therefore remain difficult to investigate. In *Gmnc* mutants however, the use of pan-CRs markers such as *Reln*, *Nhlh2* or *Tbr1* allows one to monitor presumed VP-derived CRs. We found these cells remain in the lateral cortex in *Gmnc* mutants, suggesting they are unable to redistribute. This is somehow surprising as examples of CRs redistribution (physiological or experimentally induced) were reported for distinct subtypes and time points (Barber et al., 2015; de Frutos et al., 2016; Griveau et al., 2010). This suggests that contact-mediated repulsion between CRs (Villar-Cerviño et al., 2013) is not the only mechanism involved in their positioning, but that local or diffusible cues also confine them to specific regions or prevent their spread in adjacent ones. Alternatively, CRs redistribution could be temporally restricted, occurring only at early stages of corticogenesis, at least for some subtypes. Another interesting observation we could make is that VP-derived CRs seem to disappear earlier than their p73^+^ counterparts. This is consistent with the hypothesis we previously proposed that CRs are fated to die and that components of their differentiation program are required to maintain them alive (Causeret et al., 2018; Moreau et al., 2023). According to this model, the lack of expression of the *Gmnc*/*Trp73* gene regulatory network would render VP-derived CRs more prone to early demise than other subtypes.

### Limitations of the study

One technical limitation of our study was the incomplete efficiency of conditional *Reln* deletion in P73^+^ CRs that prevented us to be fully conclusive regarding Reln-dependent and - independent functions of CRs. We previously reported a similar issue attempting to conditionally inactivate *Gmnc* in CRs (Moreau et al., 2023) raising the possibility that CRs are somehow less sensitive to Cre-mediated recombination than other cell types. Another limitation was the hydrocephaly that progressively develops in *Gmnc* mutants at postnatal stages that prevented us from investigating cortical morphogenesis at later stages, when cortical lamination and hippocampal morphogenesis are fully achieved. We believe that the increased intracranial pressure and progressive mortality associated with hydrocephaly (Terré et al., 2016) would have hindered correct interpretation of the data. Finally, our work opens a number of questions that could not be answered in this study: What are the mechanisms allowing the residual production of p73^+^ CRs in *Gmnc* mutants? What are the molecular players behind the Reln-independent function of CRs in hippocampal morphogenesis? Future work will be required to address these points.

## Supplementary Figure legends

**Figure S1. Expression of *Nhlh2*.** (A-C) *In situ* hybridization for *Nhlh2* at E12 and E13 (A), and at E16 in the lateral cortex (B) or hippocampus (C) in control and *Gmnc^-/-^* embryos. The presence/absence of *Nhlh2*^+^ cells in the MZ is indicated by filled/empty arrowheads, respectively. Scale bars: 200µm.

**Figure S2. Reln loss in *Gmnc* mutants.** Immunostaining for Reln (red) along the rostro-caudal axis of the brain in control and *Gmnc^-/-^* animals at P1. Scale bar: 500µm.

**Figure S3. Layering of the dorsal cortex in *Gmnc* mutants.** (A) Quantification showing that superficial layers 2-4 are thinner in *Gmnc* mutants compared to controls whereas deeper layers 5 or 6 are unaffected. * p<0.0001 using Mann Whitney test. (B) Immunostaining for Reln (red) and Foxg1 (green) in control or *Gmnc^-/-^* E18 embryos showing MZ invasion by Foxg1^+^ neurons in mutants. (C) Immunostaining for Prox1 in the dorsal cortex of control or *Gmnc^-/-^* E18 embryos and quantification showing no defects in the number or position of Prox1^+^ interneurons. Scale bars: 100µm.

**Figure S4. CRs distribution and Reln loss in Reln cKO.** (A) Immunostaining for Tomato in control and Reln cKO showing the normal numbers and distribution of CRs. (B) Immunostaining for Reln (green) and Tomato (red) in the neocortex or hippocampus of P1 control or *Reln^cKO^* animals. (C) Immunostaining for Tomato, Reln and Foxg1 in Reln cKO showing that cortical neurons invading the MZ (dashed line) can be found in direct proximity with either Tomato^+^ or Reln^+^ cells. (D, E) Immunostaining for Reln in the lateral cortex (D) and hippocampus (E) of control and *Reln^cKO^*animals. Scale bars: 500µm in A, D, E, 50µm in B, 100µm in C.

## Acklowledgements

We are grateful to all members of the Pierani and Spassky labs for stimulating and helpful discussions and Anne Teissier for critical reading of the manuscript. We thank Marine Luka and the LabTech Single-Cell@Imagine for library preparation, Cecile Masson and all the staff from the Imagine genomic and bioinformatics core facilities as well as the Institut Français de Bioinformatique for providing access to the R-Studio service of the IFB-core cluster. We acknowledge the histology and cell imaging facilities of the Structure Fédérative de Recherche Necker (Inserm US24, CNRS UAR3633) and the NeurImag Imaging core facility of IPNP. We thank Leducq establishment for funding the Leica SP8 confocal/STED 3DX system at IPNP and Association pour la Recherche sur le Cancer for funding the Nanozoomer slide scanner at SFR Necker. We thank the Imagine Institute LEAT and IPNP animal facility for animal care, especially the zootechnicians in charge of our colonies.

## Author contribution

Conceptualization: VE, AP, NS, FC

Methodology: VE, BB, YS, PA, MXM, AP, NS, FC

Software: FC

Validation: VE, BB, YS, JSM, ED, NS, FC

Formal analysis: VE, BB, FC

Investigation: VE, BB, YS, JSM, ED, FC

Resources: RS, AP, NS, FC

Data curation: VE, FC

Writing - original draft: FC

Writing - review & editing: VE, BB, YS, ED, RS, AP, NS, FC

Visualization: VE, FC

Supervision: AP, NS, FC

Project administration: FC

Funding acquisition: NBB, AP, NS, FC

