## Supplementary figures and images for "Differential contribution of P73^+^ Cajal-Retzius cells and Reelin to cortical morphogenesis"

### Supplementary Figure 1

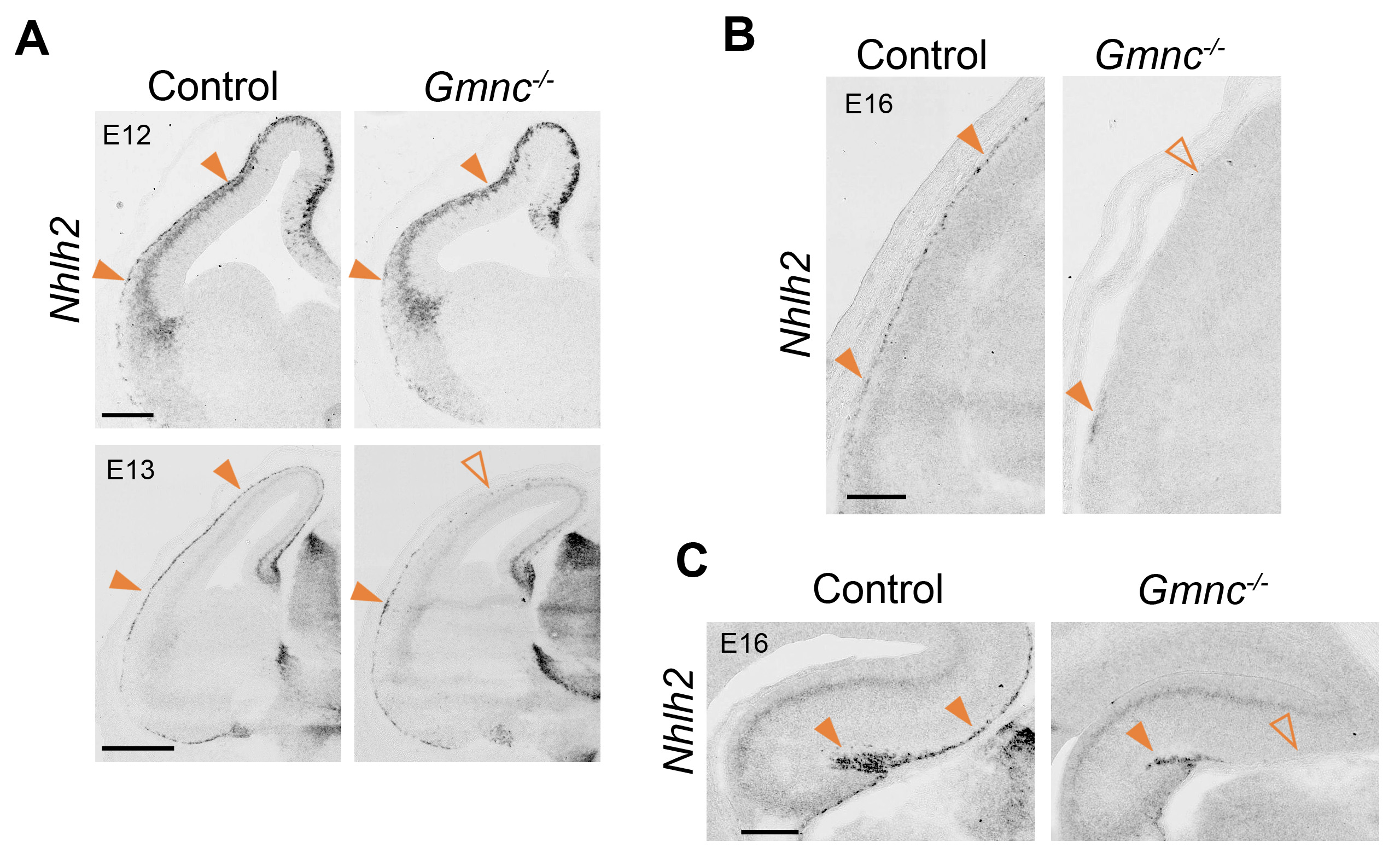

### Supplementary Figure 2

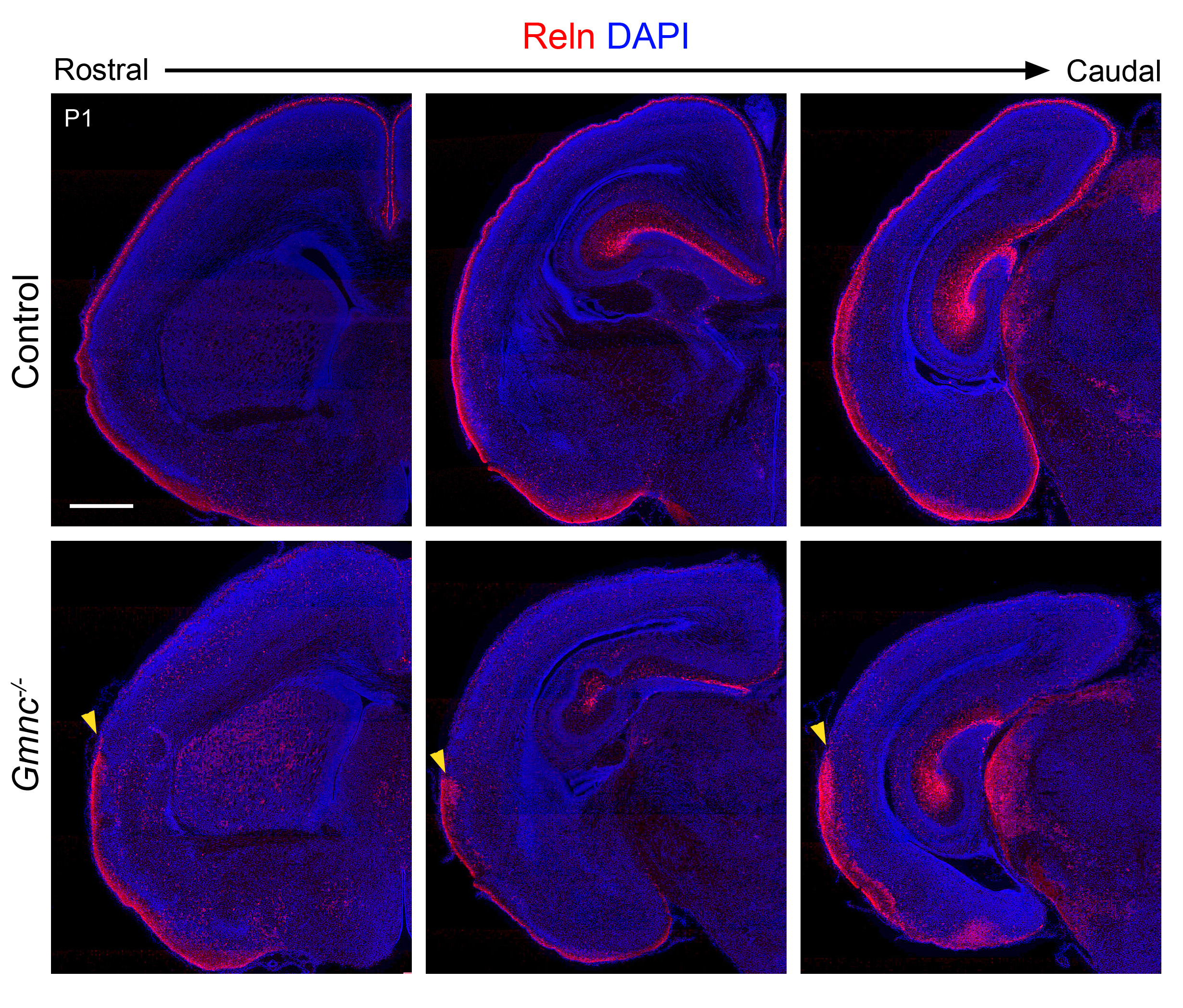

### Supplementary Figure 3

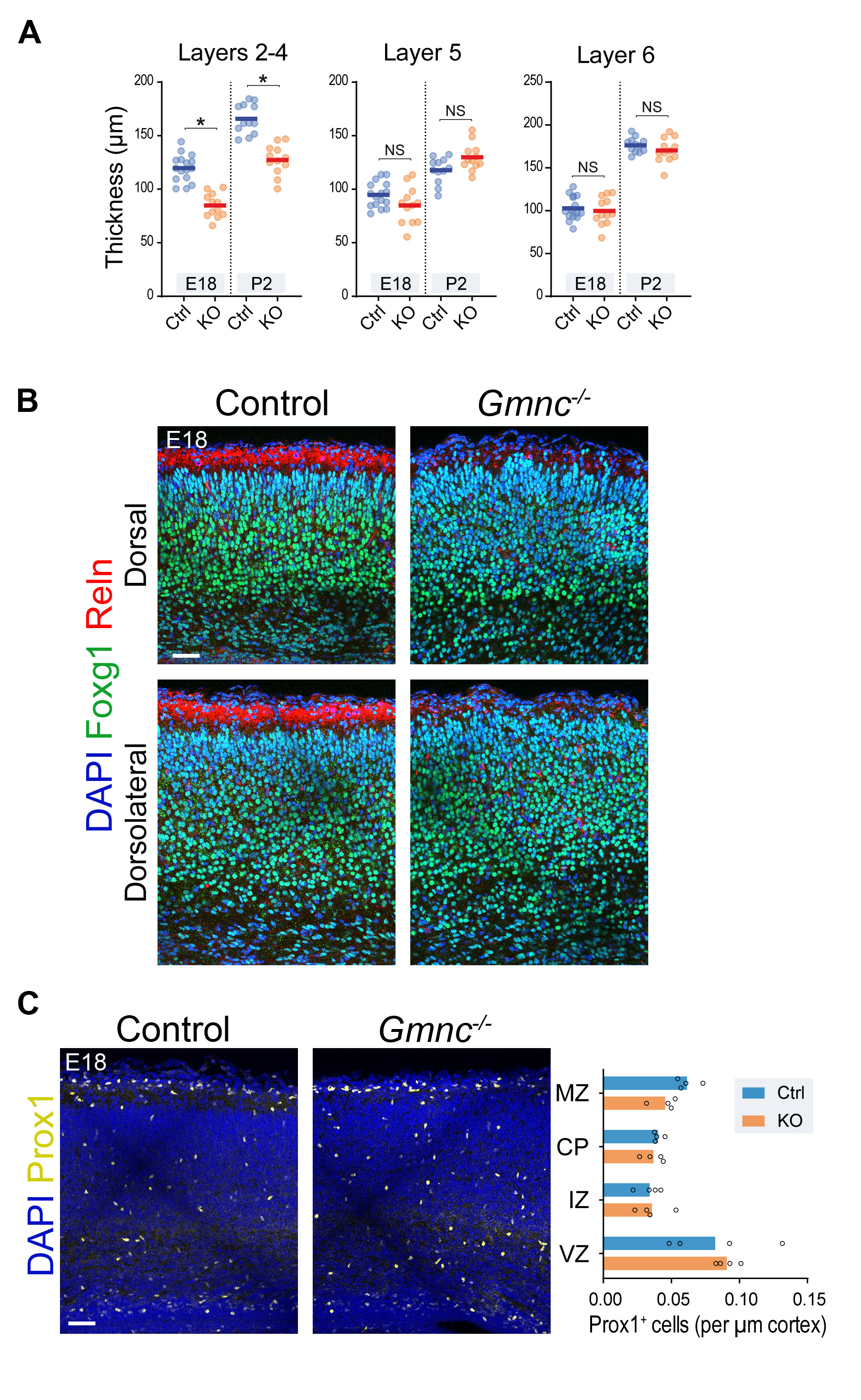

### Supplementary Figure 4

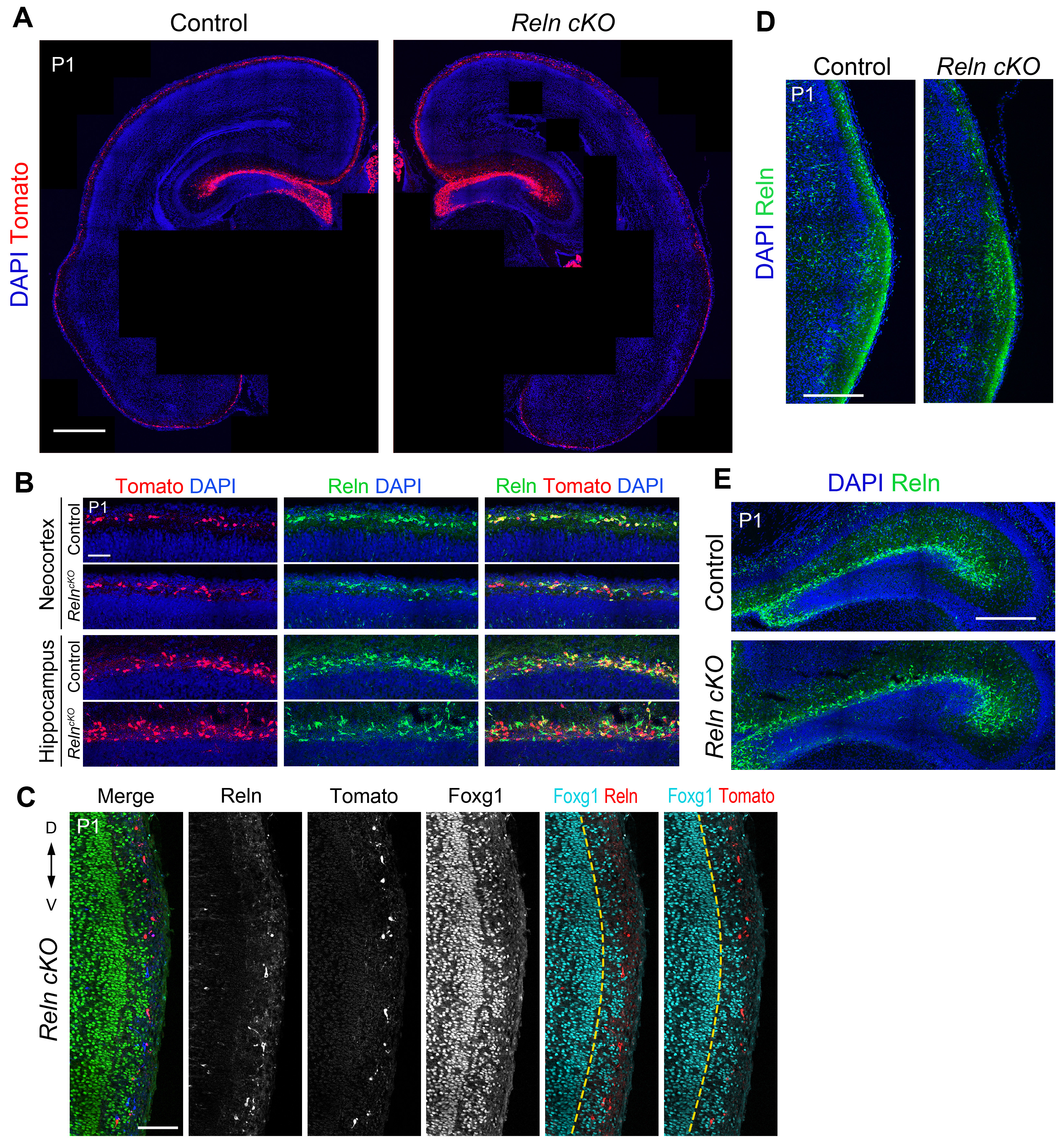
